# GROOLS: reactive graph reasoning for genome annotation through biological processes

**DOI:** 10.1101/117994

**Authors:** Jonathan Mercier, Adrien Josso, Claudine Médigue, David Vallenet

## Abstract

**Background:** High quality functional annotation is essential for understanding the phenotypic consequences encoded in a genome. Despite improvements in bioinformatics methods, millions of sequences in databanks are not assigned reliable functions. The curation of protein functions in the context of biological processes is a way to evaluate and improve their annotation.

**Results:** We developed an expert system using paraconsistent logic, named GROOLS (Genomic Rule Object-Oriented Logic System), that evaluates the completeness and the consistency of predicted functions through biological processes like metabolic pathways. Using a generic and hierarchical representation of knowledge, biological processes are modeled in a graph from which observations (i.e. predictions and expectations) are propagated by rules. At the end of the reasoning, conclusions are assigned to biological process components and highlight uncertainties and inconsistencies. Results on 14 microbial organisms are presented.

**Conclusions:** GROOLS software is designed to evaluate the overall accuracy of functional unit and pathway predictions according to organism experimental data like growth phenotypes. It assists biocurators in the functional annotation of proteins by focusing on missing or contradictory observations.

## 1 Background

Assigning functions to all predicted proteins from genome sequencing projects remains challenging despite improvements in bioinformatics methods as illustrated by the second critical assessment of functional annotation initiative (CAFA2) [1]. Millions of protein entries are thus not assigned reliable functions in databanks due to the lack of trustworthy annotations and the drawbacks of homology-based predictions [2]. Using this genomic data, genome-scale metabolic networks can be obtained and contain all of the known metabolic reactions in a given organism and the genes that encode each enzyme. High quality functional annotation is thus an essential step to correctly predict the chemical transformations that may occur in a living cell. From these networks, metabolic models can be derived to study metabolic and growth functions notably by using flux balance analysis [3]. The predictive power of these models is very sensitive to incomplete knowledge and their manual curation is time-intensive although several algorithms can be used to predict which reactions are missing by comparing *in silico* growth simulations to experimental results [4].

In order to automatically annotate protein sequences with a high degree of accuracy (i.e. by limiting annotation error propagation between homologous proteins), the UniProt consortium has developed the UniRule system [5]. It contains a set of annotation rules that are defined by biocurators using UniProtKB/Swiss-Prot annotations as a gold standard. These rules are based on the presence of specific protein signatures together with taxonomic constraint to predict biological features and functions of unreviewed proteins of the UniProtKB/TrEMBL section. The process annotates proteins independently without considering other predicted functions in the studied organism. Improving and validating functional annotation of proteins in a metabolic context was shown of interest in several microbial genome annotation systems such as SEED [6], MicroScope [7, 8] and IMG [9]. Similarly to UniRule that predicts protein functions, Genome Properties [10, 11] and IMG [12] systems have defined rules to predict biological processes. These rules are a mix of disjunctions and conjunctions of process components that are hierarchically organized to represent, for example, a pathway composed of reactions that are catalyzed by enzymes made of polypeptides in complex. Rules are evaluated using protein annotations as facts to determine if a minimal set of required functions is predicted to infer the presence or absence of a process in an organism. In the IMG system, additional rules based on biological processes are used to predict phenotypes like auxotrophy/prototrophy for biosynthetic pathways or compound utilization for degradation pathways.

Here, we present an expert system, named GROOLS (Genomic Rule Object-Oriented Logic System), that uses paraconsistent logic to assist biocurators in the evaluation of the functional annotation of genes. The GROOLS method is inspired from a previous work about the HERBS software (A. Viari *et al.*, unpublished data) that was presented in 2003 during a workshop of the HAMAP project [13]. The objective is to automate human expert reasoning applied in the curation of genome annotation. Similarly to previously mentioned systems (i.e. Genome Properties and IMG), GROOLS evaluates the completeness and the consistency of predicted functions through biological processes like metabolic pathways. Our objective was to propose a framework made of a generic data model to embrace various sources of biological process definitions. Moreover, the tool should consider potential contradictions or incomplete information and should make a clear distinction between what is predicted and what is expected during the reasoning. Expectations could be background knowledge on the organism, experimental results like growth phenotypes or simply biological hypotheses provided by users. At the end of the reasoning, conclusions should highlight confirmed/missing predictions and potential conflicts between predictions and expectations. Following a description of the GROOLS method, results on 14 microbial organisms, having available growth phenotype data, are presented and illustrated by two examples. Finally, we conclude on the GROOLS functionalities, potential applications and improvements.

## 2 Methods

### 2.1 Definitions and biological knowledge representation

GROOLS uses a generic model to represent biological processes in a hierarchical structure (Figure 1). Indeed, GROOLS knowledge representation is based on a conceptual graph [14] with two types of concepts:

- Prior Knowledge (PK) concepts represent biological processes and their components (e.g. a metabolic pathway and its reactions)
- Observation (O) concepts are information about PK concepts in a given organism.

The hierarchical organization of PK concepts is made by two types of edges in the conceptual graph:

- ‘part’ relations describe compositions (e.g. a metabolic pathway is composed of a set of reactions)
- ‘subtype’ relations describe generalizations and specializations (e.g. pathways 1.1 and 1.2 are two variants of the generalized metabolic pathway 1).

Edges are directed and labelled with the suffix “-of” (i.e. ‘part-of’ and ‘subtype-of’) when the relation is from the parent concept to the child concept and, conversely, with the prefix “has-” (i.e. ‘has-part’ and ‘has-subtype’).

To give clues about the existence of PK concepts in a given organism, three types of Observation are used in GROOLS:

- Computations are bioinformatics predictions (e.g. genome annotation methods that predict protein gene functions like enzymatic activities and their corresponding catalyzed reactions)
- Experimentations are empirical evidences (e.g. growth phenotypes with defined metabolites as sole carbon and energy source)
- Curations are human expertises supported by direct or indirect evidences (e.g. the validation of a bioinformatics prediction with additional literature evidences)

Finally, Observations are linked to PK concepts with two types of relations:

- ‘prediction-of’ edges for Computation and Curation observations
- ‘expectation-of’ edges for Experimentation and Curation observations.

This GROOLS conceptual graph model is generic enough to capture the overall background knowledge on biological processes represented by PK concepts (e.g. metabolic pathways from all domains of life) and organism specific knowledge represented by Observations that come from experimentations or its genome annotation. Each observation corresponds either to an assertion or a rejection of a PK concept. Such states are represented by the true (t) and false (f) values also named truth values in classical logic. Observations will be propagated through the conceptual graph to assign PK concepts prediction or expectation values (see the “Reactive reasoning” paragraph). These values answer the following questions: “Is the PK concept predicted in the studied genome?” and “Is it expected in the organism?”.

### 2.2 Observation and Prior Knowledge truth value powersets

In biology, several experimentations or bioinformatics predictions may be related to a same PK concept. To capture facts from this observable space, Observations are grouped by their PK target and relation type (i.e. ‘prediction-of’ or ‘expectation-of’) resulting from an Observation truth value powerset of order 2 (ℙ(2)). These Observations truth value sets (Otvs) can take the following values: {t} (true-only Observations), {f} (false-only Observations), {t,f} (true and false Observations) or 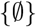 (no Observation). To rank Otvs according to their level of truth or falsehood, they are associated to the truth degrees 1, 0, 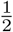, 0 and to the falsehood degrees 0, 1, 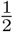, 0 for the {t}, {f}, {t,f} and 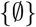 sets, respectively.

**Figure 1.**
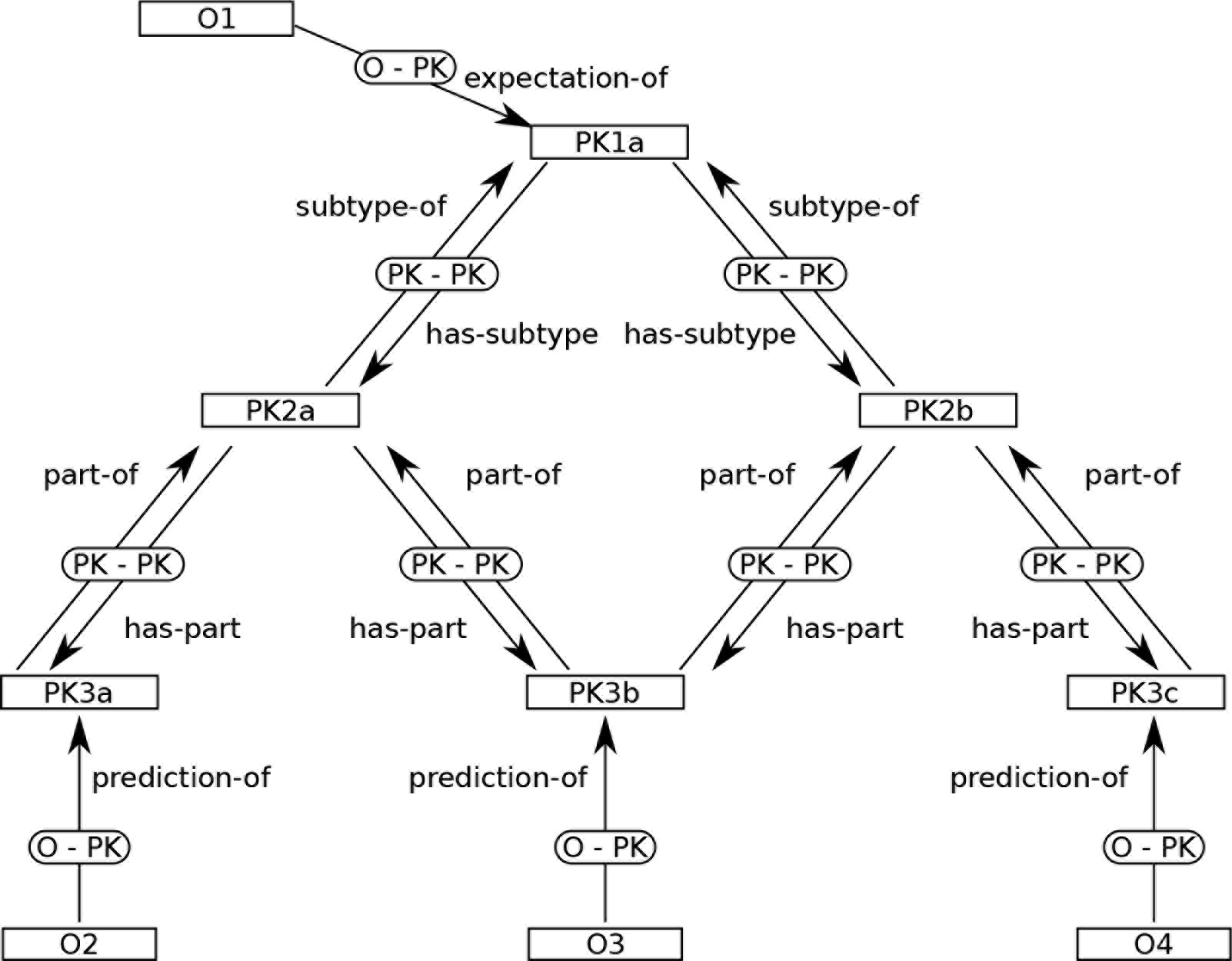
Illustration of the GROOLS conceptual graph model. This figure gives an illustration of the GROOLS generic model that is used to represent any biological processes. The model is a conceptual graph with two types of concepts (rectangle nodes): Prior Knowledge (labelled PK) and Observation (labelled O). Relations between concepts are represented by edges named PK-PK (between two PK concepts) or O-PK (between O and PK concepts). Four types of relations are available in the model (i.e. ‘subtype’, ‘part’, ‘expectation’ and ‘prediction’) and are labelled with the ‘-of’ suffix or the ‘has-’ prefix according to the edge direction.

In the conceptual graph, Observation sets are propagated directly to PK but also indirectly through PK concept relations. Thus, both prediction and expectation values of PK hold sets of direct and indirect Observations leading to a powerset of order 4 (i.e. ℙ(4) corresponding to 16 sets listed in Table 1). Actually, GROOLS is based on a paraconsistent logic that uses an ensemblist representation of true and false value combination. It allows us to deal, without approximation, with uncertainty or contradiction for PK predictions and expectations. An algebraic structure behind these 16 subsets was proposed by Shramko *et al.* [15, 16] and is known as a trilattice SIXTEEN_3_ with the following distinct partial orders: the truth, the falsehood and the degree of information. In GROOLS, the degree of truth/falsehood of each subset is computed by the sum of Otvs truth/false degrees present in the powerset divided by the degree of information (Table 1). Finally, to determine the degree of belief, PK predictions and expectations are ranked by decreasing truth degree, then by increasing degree of falsehood and information, in case of equality (Table 1).

**Table 1.**
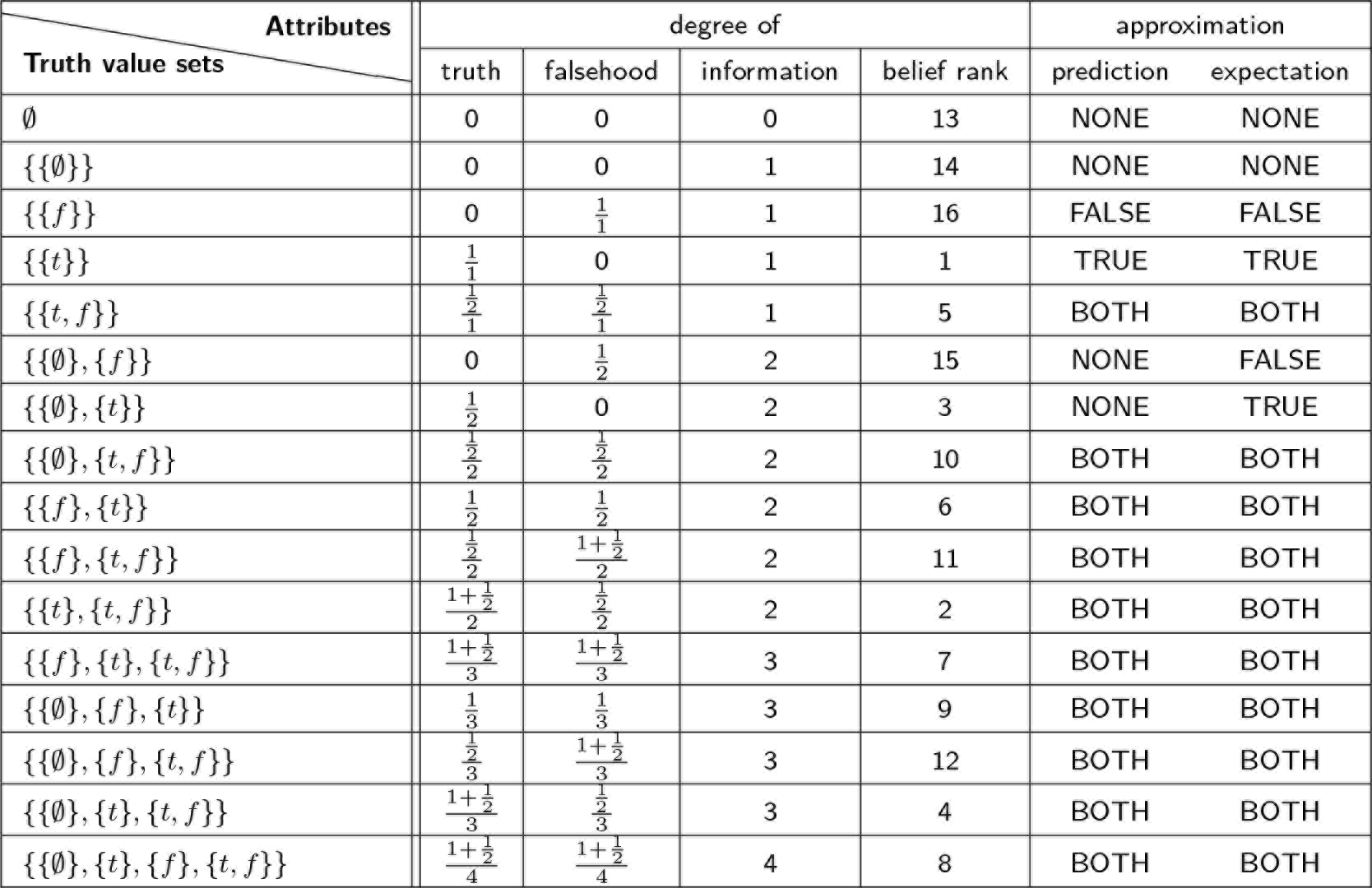
The sixteen truth value sets and their attributes

### 2.3 Reactive reasoning

The GROOLS reasoner propagates information from Observation to PK concepts and between PK using inference rules in three steps: propagation of predictions, propagation of expectations and PK conclusions. Thanks to the reactive nature of GROOLS reasoning, the declaration of new Observations can be made at any time and the logical consequences are then propagated locally in the graph without re-evaluating all PK predictions and expectations. A graphical illustration of GROOLS reasoning is given in Additional file 1.

#### 2.3.1 Propagation of predictions

PK prediction values are assigned by collecting truth value sets from direct Observations (Computation or Curation type) and child PK predictions. The algorithm propagates predictions from the leaves to the roots of the conceptual graph using two distinct rules: one for ‘has-part’ relations (Rule 1) and another for ‘has-subtype’ relations (Rule 2). For the Rule 1, PK predictions are the union of Otvs (Observation truth value sets) and Ptvs (Prediction truth value sets) from child PK concepts. In Rule 2, PK predictions are the union of Otvs and Ptvs from child PK concepts having the greatest degree of belief (i.e. the smallest belief rank).

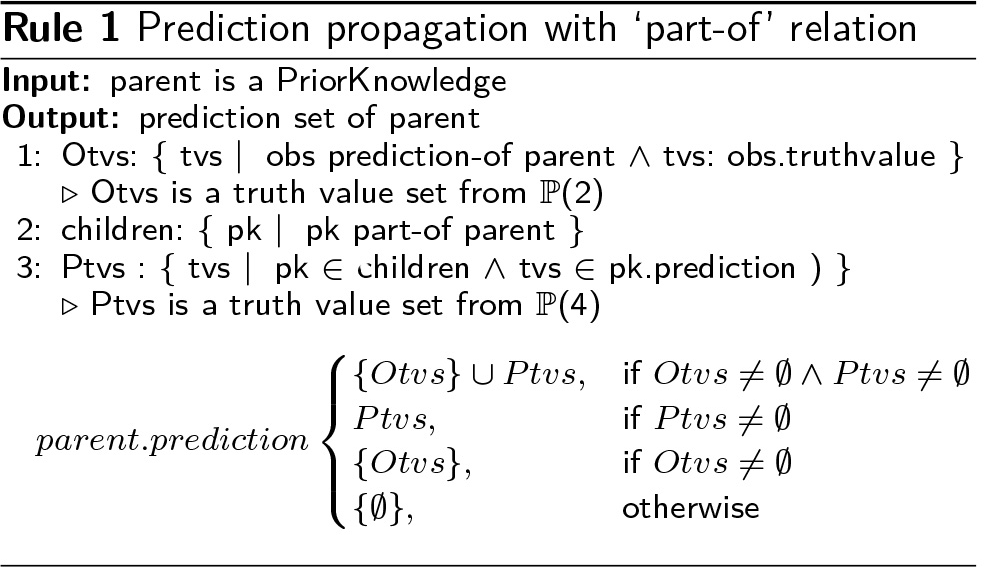

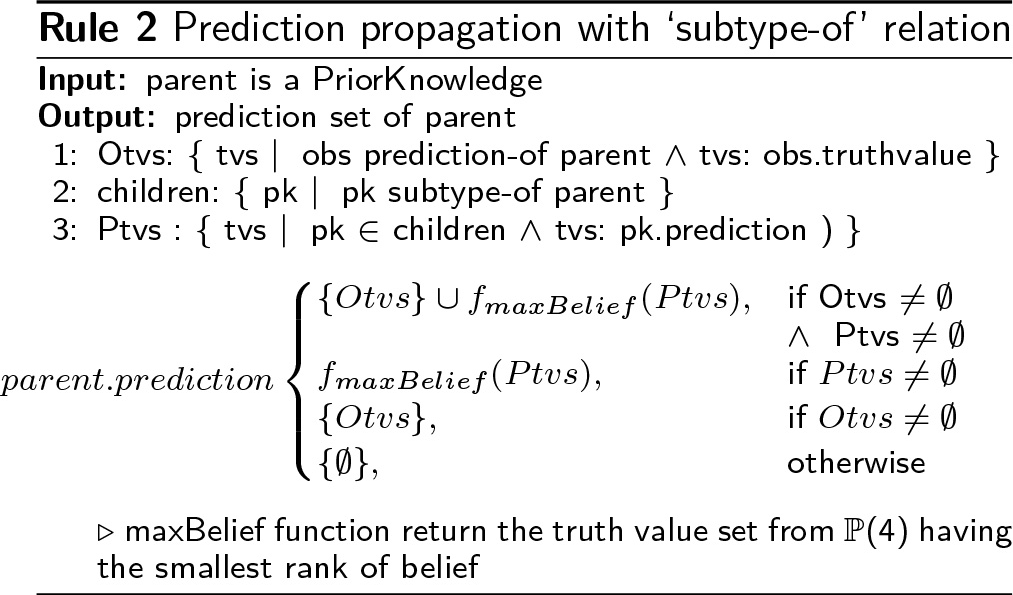

#### 2.3.2 Propagation of expectations

In a second step, PK expectations are propagated from the roots to the leaves of the graph by collecting truth value sets from direct or indirect Observations (Experimentation or Curation type). For the ‘has-part’ relations, PK expectations are the union of Otvs and Etvs (Expectation truth value sets) from parent PK concepts (Rule 3). For the ‘has-subtype’ relations, parent PK expectations are propagated to child PK that have the greatest degree of belief for their prediction with the exception of false-only expectations ({{f}}) that are propagated to all child PK (Rule 4).

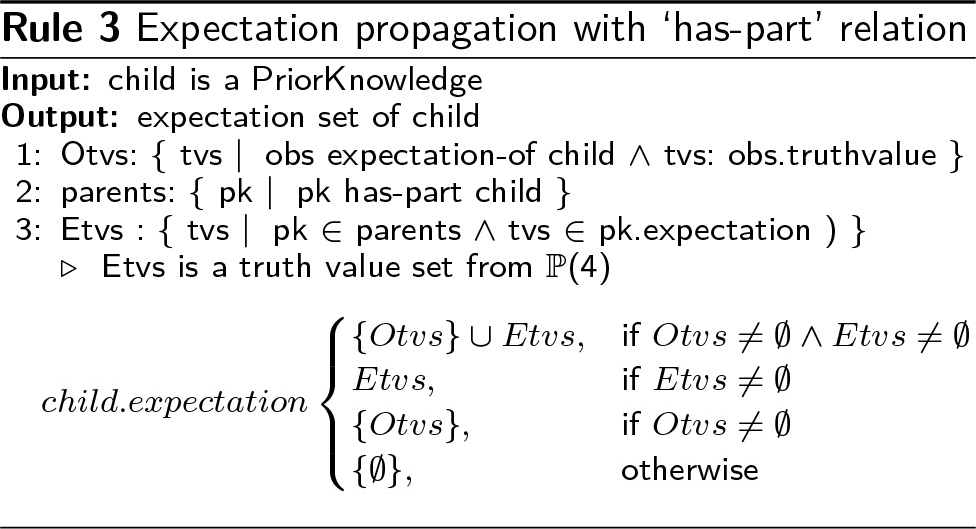

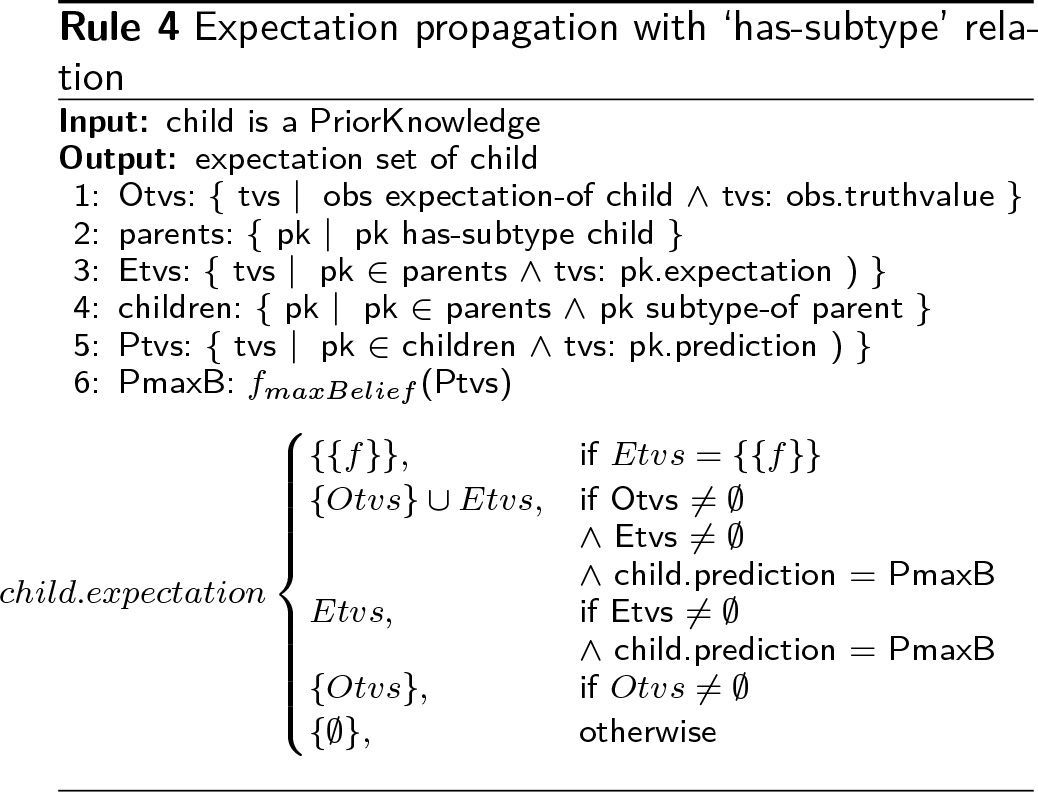

#### 2.3.3 PK conclusions

At the end of the reasoning, to ease human interpretation of PK predictions and expectations, truth value sets are approximated in four values: TRUE, FALSE, BOTH, NONE (Table 1). When both true and false values are present, the approximated value is BOTH to express contradiction. For predictions, if the set contains an unknown value 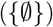 and a true or false value, the approximated value is NONE meaning that some parts of the concept are unknown. For expectations, unknown values are not considered in the approximation as they do not represent missing part of a concept but lack of information about a higher concept among others having empirical evidences (true or false expectation values). Thus 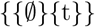 and 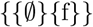 expectation sets are approximated to TRUE and FALSE, respectively.

The combination of these approximated values of PK predictions and expectations lead to 16 different states (Table 2). They express a conclusion about the presence/absence of a concept in an organism according to experiments and genome annotations and point out potential contradictions and uncertainties. For examples, a “Confirmed Presence” is used when the concept is both predicted and expected. An “Unexpected Presence” corresponds to a predicted process for which direct or indirect experimental evidences indicate that it does not occur in the studied organism. If experiments contradict themselves (BOTH expectation) while the process is predicted, the conclusion is an “Ambiguous Presence”. Finally, as a last example, the “Missing” state is an absence of prediction (NONE prediction) while the process is experimentally observed. In this case, the genome annotation should be improved to find candidate proteins for the missing functions.

**Table 2.** Conclusion table

| Prediction<br>Expectation | TRUE | FALSE | BOTH | NONE |
| --- | --- | --- | --- | --- |
| TRUE | Confirmed P. | Unexpected A. | Contradictory A. | Missing |
| FALSE | Unexpected P. | Confirmed A. | Contradictory P. | Absent |
| BOTH | Ambiguous P. | Ambiguous A. | Ambiguous C. | Ambiguous |
| NONE | Unconfirmed P. | Unconfirmed A. | Unconfirmed C. | Unexplained |

P.: Presence, A.: Absence, C.: Contradiction

#### 2.3.4 Dispensable, falsehood and specific modes

Three additional reasoning modes have been developed to handle particular cases in the reasoning. In the Dispensable mode, PK concepts can be flagged as nonessential meaning that their presence is not required to realize a biological process. In this case, predictions and expectations are not propagated to PK concepts labelled as “Dispensable”. In the Falsehood mode, leaf PK concepts (i.e. concepts without ‘has-part’ and ‘has-subtype’ relations) with no direct Observations have their prediction assigned to false-only ({{f}}). This reasoning mode assumes that all leaf concepts of the graph are predictable, and thus, an absence of Observation corresponds to a false prediction. In the Specific mode, a PK is labelled as “Specific” when it is linked to only one parent PK by a ‘part-of’ relation. Moreover, in this mode, Rule 1 (propagation of child PK predictions to parent PK using ‘part’ relations) is adapted: child PK with unknown prediction 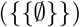 are not considered when at least one other child PK has a true-only prediction ({{t}}). Using this Specific mode, reasoning is less strict as partial evidences on child PK are sufficient to predict a parent PK.

### 2.4 Biological data

Prior Knowledge (PK) concepts represent biological processes and their components (e.g. a metabolic pathway and its reactions). Genome Properties [10] and UniPathway [17] resources are currently supported by GROOLS to reconstruct two distinct conceptual graphs.

From the Genome Properties description file (version 3.2, GenProp_3.2_release.RDF), “Properties”, “Components” and “Evidences” were extracted and transformed in PK concepts. ‘Part’ relations were established between “Properties” and “Components”, and ‘subtype’ relations between “Components” and “Evidences”. In the Genome Properties model, “Evidences” correspond either to pathway variants or predictors (i.e. leaf concepts in the graph). Predictors are protein family Hidden Markov Models (HMMs) from PFAM [18] and TIGRFAM [11] databases. Several leaf “Evidences” linked to a “Component” represent distinct protein families performing the same function (e.g. isoenzymes).

From the UniPathway Open Biological Ontology (OBO) file, UPA (pathways), ULS (linear sub-pathways), UER (enzymatic reactions) and UCR (chemical reactions) were extracted and transformed in PK concepts. For ULS linked to a same UPA, additional concepts, named Variants, were added when several paths of ULS, that share input/output compounds, exist. Other Variants were made for UER that are specializations of generic UER (i.e. “has_alternate_enzymatic_reaction” relations in the OBO file). According to “is_a” and “super_pathway” relations in the OBO file, UPA were linked together by ‘subtype’ relations in the conceptual graph. ‘Subtype’ relations were also used to link Variants to UPA or UER. Other relations between PK concepts were converted to ‘part’ relations.

To populate Genome Properties conceptual graph, Observations of Computation type can be extracted from UniProtKB database [5] in which each protein entry is linked to predicted PFAM [18] and TIGRFAM [11] families. For UniPathway, UniProtKB annotations can be used and inserted in the conceptual graph as Observations of UER (UniPathway Enzymatic Reactions) using their identifiers (when provided in the protein entry) or their cross-referenced EC (Enzyme Commision) numbers. Annotations from the MicroScope platform [8] can be used as a second resource of UniPathway Observations. MicroScope contains automatic and curated annotations of microbial genomes. Similarly to UniProtKB, protein annotated with EC numbers are converted in UER Observations. In addition, Observations on UCR (UniPathway Chemical Reactions) are made using cross-referenced MetaCyc [19] or RHEA [20] reactions that are annotated in MicroScope protein entries. Automatic annotations from MicroScope are considered as Observations of Computation type, whereas curated annotations are considered as Observations of Curation type (i.e. Observation truth value sets are propagated both on PK predictions and expectations). A specific case concerns MicroScope pseudogenes which predicted original functions are considered as false Observations of UCRs and UERs (i.e. corresponding reactions should not occur anymore in the organism as pseudogenes do not produce functional proteins).

GROOLS can integrate Phenotype MicroArrays (PMs) BIOLOG experiments as Observations of Experimentation type. This quantitative growth phenotype data should be first discretized using the omp R package [21] with “grofit” aggregation method and weak discretization (-a, -w and -z options of run_opm.R program). Using Observations with a true (growth) or false (no growth) truth value, the results obtained with carbon and nitrogen sources (PM1, 2 and 3 plates) are then linked to their corresponding Genome Properties (16 pathways) or UniPathway (37 pathways) PK concepts (i.e. the metabolic pathway for the degradation of the tested compound). For ambiguous discretizations (NA values), both true and false Observation concepts are created. Regarding prototrophic organisms, true Observations on amino acid biosynthesis pathways can be added as a second source of Experimentation corresponding to 15 pathways in Genome Properties and 20 in UniPathway.

### 2.5 Statistics

Statistics were computed to evaluate predictions in confrontation to biological expectations. In this evaluation, only concepts with univocal expectations ({{t}} and {{f}} values) are considered. Concepts are classified using their conclusion value as follow:

- True Positive (TP): ‘Confirmed Presence’
- True Negative (TN): ‘Confirmed Absence’ and ‘Absent’
- False Positive (FP): ‘Unexpected Presence’ and ‘Contradictory Presence’
- False Negative (FN): ‘Missing’, ‘Unexpected Absence’ and ‘Contradictory Absence’ concepts.

Accuracy (ACC) is computed as follow:

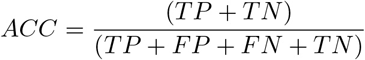

### 2.6 Implementation

GROOLS is built on object-oriented logic programming. This paradigm facilitates the representation of biological knowledge with Prior Knowledge concepts and their relations. A first prototype of GROOLS (https://github.com/Grools/grools-drools-checker) was written using the DROOLS framework (https://www.drools.org, a Business Rules Management System based on an object-oriented rule language). In the present version of GROOLS software (https://github.com/Grools/grools-application), rules were directly implemented in Java with functional programming to tackle performance issues during the graph reasoning. The reasoner is available in a separate library (https://github.com/Grools/grools-reasoner).

GROOLS contains plugins to construct the PK graph from two databases: UniPathway (in OBO format) and Genome Properties (in non-standard format). Using this plugin system, database file content can be improved by users. Moreover, other resources on biological processes can be integrated without changing the reasoning library.

For Observations, an unified CSV file format is used to represent Computation, Curation and Experimentation data. Each row is composed of seven fields to describe an Observation: “Name”, “EvidenceFor” (name of the related Prior Knowledge), “Type” (Computation, Curation or Experimentation), “isPresent” (true or false), “Source” (e.g. UniProtKB, MicroScope, BIOLOG), “Label” and “Description”.

Three shell scripts are provided as example to run GROOLS on Genome Properties using UniProtKB proteomes (“uniprot_genpropToGrools.sh” script) and on UniPathway using MicroScope or UniProtKB annotations (“microscope_upaToGrools.sh” and “uniprot_upaToGrools.sh”, respectively).

GROOLS results are provided in CSV format and in HTML pages containing graphical representations of the conceptual graph for each biological process. GROOLS execution time is about 1 to 2 minutes to completely analyze a genome. The reasoning part takes only few seconds while most of the execution time is spent in data extraction and graphical output generation.

## 3 Results

### 3.1 GROOLS reasoning test case

As a test case, GROOLS was executed on 14 organisms for which UniProtKB and MicroScope annotations are available. The reasoning was made on Genome Properties and UniPathway conceptual graphs, enabling or not the falsehood (for Genome Properties) and specific (for UniPathway) modes. In addition, the dispensable mode was enabled for Genome Properties as this resource contains information about non-essential components of biological processes. For Observations of Experimentation type, growth phenotypes (PM BIOLOG technology) were retrieved for these organisms on the Microme project website (http://www.microme.eu) and linked to their corresponding degradation pathway PK concepts. Given that studied organisms are prototroph, additional Observations with true truth value were added on amino acid biosynthesis pathway concepts (i.e. these pathways are expected in the organisms). Detailed results on the 14 studied organisms are available at this Web address: https://www.genoscope.cns.fr/agc/grools. Statistics on GROOLS conclusions are reported in Table 3. Accuracies of pathway and functional unit predictions were computed only for pathways with univocalexpectations (i.e. pathway concepts with true-only or false-only Observations of Experimentation type).

**Figure 2.**
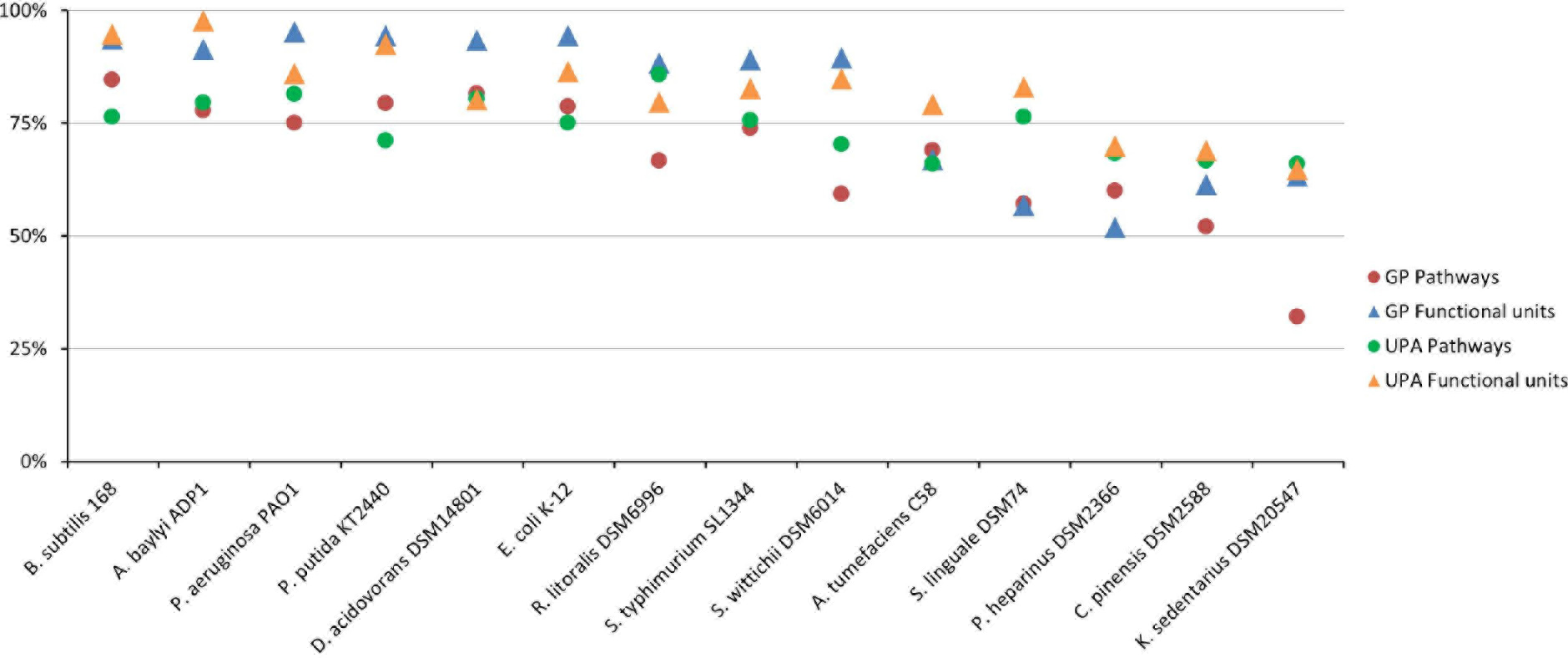
Evaluation of prediction accuracy. Accuracy for pathways and functional unit predictions was evaluated for Genome Properties (GP) and UniPathway (UPA) in 14 organisms. For Genome Properties, GROOLS falsehood mode is activated. For UniPathway, the specific mode is activated and annotations are from MicroScope.

**Table 3.** GROOLS statistics on pathways and functional units for Genome Properties and UniPathway. Statistics are computed for the 14 studied organisms.

|  |  | Genome Properties |  | UniPathway |  |  |  |
| --- | --- | --- | --- | --- | --- | --- | --- |
|  |  | UniProtKB | UniProtKB falsehood | UniProtKB | MicroScope | UniProtKB specific | MicroScope specific |
| **Pathway conclusions** | **Exp. / Pred.** |  |  |  |  |  |  |
| Confirmed Presence | TRUE / TRUE | 195 | 195 | 241 | 277 | 339 | 393 |
| Unexpected Absence | TRUE / FALSE | 0 | 12 | 0 | 0 | 0 | 0 |
| Contradictory Absence | TRUE / BOTH | 0 | 56 | 0 | 5 | 0 | 5 |
| Missing | TRUE / NONE | 68 | 0 | 232 | 191 | 137 | 80 |
| Unexpected Presence | FALSE / TRUE | 13 | 13 | 33 | 50 | 79 | 121 |
| Confirmed Absence | FALSE / FALSE | 0 | 66 | 0 | 0 | 0 | 0 |
| Contradictory Presence | FALSE / BOTH | 0 | 40 | 0 | 0 | 0 | 0 |
| Absent | FALSE / NONE | 106 | 0 | 275 | 258 | 229 | 187 |
| Ambiguous Presence | BOTH / TRUE | 15 | 15 | 47 | 68 | 121 | 178 |
| Ambiguous Absence | BOTH / FALSE | 0 | 13 | 0 | 0 | 0 | 0 |
| Ambiguous Contradiction | BOTH / BOTH | 0 | 24 | 0 | 3 | 0 | 3 |
| Ambiguous | BOTH / NONE | 37 | 0 | 243 | 218 | 172 | 120 |
| Unconfirmed Presence | NONE / TRUE | 1222 | 1222 | 728 | 946 | 1130 | 1409 |
| Unconfirmed Absence | NONE / FALSE | 0 | 3453 | 0 | 0 | 0 | 0 |
| Unconfirmed Contradiction | NONE / BOTH | 0 | 726 | 0 | 5 | 0 | 4 |
| Unexplained | NONE / NONE | 4238 | 59 | 2359 | 2137 | 1951 | 1658 |
| **Pathway statistics** |  |  |  |  |  |  |  |
| True Positive |  | 184 | 184 | 200 | 238 | 272 | 310 |
| True Negative |  | 106 | 66 | 188 | 176 | 159 | 130 |
| False Positive |  | 13 | 53 | 19 | 31 | 48 | 77 |
| False Negative |  | 65 | 65 | 190 | 152 | 118 | 80 |
| Accuracy |  | 78.80% | 67.93% | 64.99% | 69.35% | 72.19% | 73.70% |
| **Functional unit statistics** |  |  |  |  |  |  |  |
| True Positive |  | 1236 | 1236 | 969 | 1719 | 982 | 1742 |
| True Negative |  | 710 | 710 | 568 | 505 | 568 | 504 |
| False Positive |  | 117 | 117 | 100 | 139 | 100 | 140 |
| False Negative |  | 396 | 396 | 511 | 342 | 465 | 283 |
| Accuracy |  | 79.14% | 79.14% | 71.55% | 82.22% | 73.29% | 84.15% |

For Genome Properties, the evaluation of functional unit predictions in confrontation to pathway expectations (i.e. from BIOLOG data and amino acid biosynthesis pathways) gives high level of accuracy (79.14%), reflecting the exquisitely sensitive searches of TIGRFAM models. In the falsehood mode, pathway prediction accuracy decreases from 78.80% to 67.93%. As expected, we observe that contradictions on pathway predictions (i.e. conclusions with BOTH predictions) are only detected using the falsehood mode. In this mode, the prediction value of a leaf component (i.e. a functional unit) is set to false-only when any corresponding PFAM or TIGRFAM domain is detected among all organism proteins. Through the propagation of these false values, pathway predictions change then from NONE to FALSE or BOTH values. One advantage of the falsehood mode is thus to raise contradictions between predictions and expectations at the pathway level (i.e. ‘Contradictory Absence’ and ‘Contradictory Presence’ conclusions). Furthermore, using falsehood mode, absence of pathways and prediction contradictions can be detected even without expectations (i.e. ‘Unconfirmed Absence’ and ‘Unconfirmed Contradiction’ conclusions).

Using UniPathway as a source of biological processes, prediction accuracies for pathways and functional units are higher for MicroScope than for UniProtKB: about twice as many functional units are correctly predicted using MicroScope annotations. Indeed, MicroScope contains further annotations on enzymes. In addition to EC numbers, reactions identifiers from MetaCyc or RHEA are generally provided in MicroScope, which facilitate the mapping of Observations on UniPathway chemical reactions (UCRs). Moreover, the UniRule annotation system is still limited to few UniProtKB entries and many entries contain original submitted annotations that are very heterogeneous in quality. Beside, the specific mode shows a positive impact on prediction accuracy. For MicroScope annotations, pathway and functional unit accuracy values increase from 69.35% to 73.70% and from 82.22% to 84.15%, respectively. Indeed, the number of ‘Missing’ conclusions (i.e. false negative predictions) is reduced to half using the specific mode. In this mode, a pathway variant is assigned a true prediction if at least one specific reaction is predicted (i.e. it does not require all reactions to be predicted). Thus, the specific mode helps GROOLS reasoning to select the correct pathway variant (i.e. the one with the greatest degree of belief) for which pathway expectations are propagated.

For BIOLOG results, it should be noticed that 23% and 37% of Genome Properties and UniPathway pathways have contradictory expectations due to ambiguous discretizations or discordant experimental replicates.

**Figure 3.**
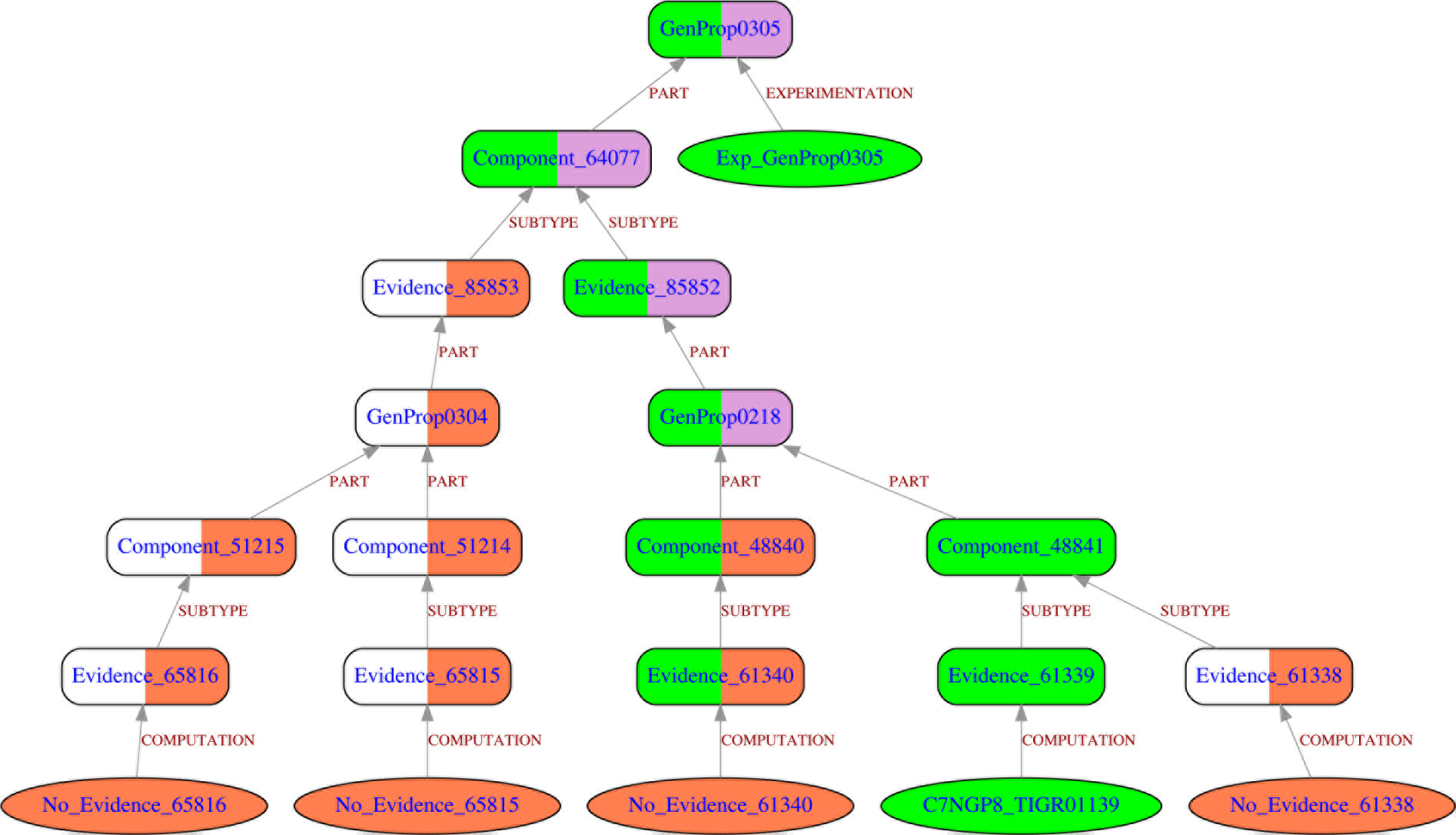
GROOLS results for cysteine biosynthesis in *Kytococcus sedentarius*. The cysteine biosynthesis in *Kytococcus sedentarius* was evaluated using Genome Properties and falsehood mode reasoning. Rounded boxes are Prior Knowledge concepts and ovals are Observations. Edges between Observations and Prior Knowledge concepts are labelled by the type of Observation whereas, between Prior Knowledge concepts, labels correspond to relation types. The color code corresponds to TRUE (green), FALSE (red), BOTH (purple) and NONE (white) values. For Prior Knowledge concepts, the colored left part corresponds to expectation value and the right part to prediction value. An interactive view of this figure is available at http://www.genoscope.cns.fr/agc/grools/paper/fig3.html

By taking into account contradictory experimental results, GROOLS may help biologists to deal with these inconsistencies by pointing out, for example, pathways with ‘Ambiguous Presence’ conclusion, which suggests that, since the pathway is predicted, the organism should be able to degrade the tested compound.

Detailed statistics on prediction accuracy for the 14 studied organisms are represented in Figure 2 (see Additional file 2 for detailed counts). Results show as expected that model bacteria (*Escherichia coli* and *Bacillus subtilis*) or other well-studied organisms (*Acinetobacter baylyi*, *Pseudomonas putida* and *P. aeruginosa*) have high prediction accuracies for functional units (accuracy > 85%) and pathways (accuracy > 70%). After manual inspection of some results, many discrepancies were found to be annotation mapping problems with UniPathway reactions (i.e. EC numbers or Meta-Cyc/RHEA reactions are not correctly cross-referenced) or incomplete pathway definition (i.e. missing variant) as illustrated in the second example below. Other organisms, like *Chitinophaga pinensis* and *Kytococcus sedentarius*, have low prediction accuracies (< 70%). These organisms are from phylum (i.e. Bacteroidetes and Actinobacteria) with limited metabolic knowledge. Potential new pathways and enzymes remain to be discovered or, more easily, predictors like TIGRFAM models could be improved to gain sensitivity as illustrated in the first example just below.

### 3.2 Genome Properties falsehood mode example

As depicted in Figure 3, cysteine biosynthesis pathway (GenProp0305) in *Kytococcus sedentarius* is flagged by GROOLS as ‘Contradictory Absence’ conclusion meaning that, although the pathway is expected in this organism, some predictions are in contradiction. Two pathway variants for cysteine biosynthesis are described in the Genome Properties model: GenProp0304 (cysteine biosynthesis, tRNA-dependent) and GenProp0218 (cysteine biosynthesis from serine). Among all functional units, only one is predicted in *K. sedentarius* (Evidence_61339 for TIGR01139 HMM) others are absent and have their prediction value set to false (falsehood mode). Rule 1 and Rule 2 of GROOLS reasoning were then applied to propagate predictions to parent concepts leading to ‘false-only’ (FALSE) and ‘true and false’ (BOTH) predictions for GenProp0304 and GenProp0218, respectively. Using rules 3 and 4, the Observation Exp_GenProp0305 (i.e. cysteine biosynthesis is expected in the organism) was propagated only through GenProp0218 as it has the greatest degree of belief for its prediction (i.e. GenProp0218 prediction value is {{t,f}} whereas GenProp0304 is {{f}}). This reasoning decision confirmed the presence of Evidence_61339 (TIGR01139) whereas Evidence_61340 (TIGR01172) is flagged as ‘Unexpected Absence’: any protein in *K. sedentarius* has significant alignment with TIGR01172 HMM. Such conclusion suggests that this missing function should be present in *K. sedentarius*. A quick investigation of TIGRFAM results showed that Ksed_26170 protein (C7NGP7 UniProtKB entry) shares an alignment with TIGR01172 HMM with a score of 200.3 that is just below the trusted cutoff (210.65). Thus, this protein is certainly a good candidate for the missing function and the TIGRFAM HMM should be updated accordingly to gain sensitivity in the detection of proteins from less-studied organisms like the actinobacteria *K. sedentarius*.

**Figure 4.**
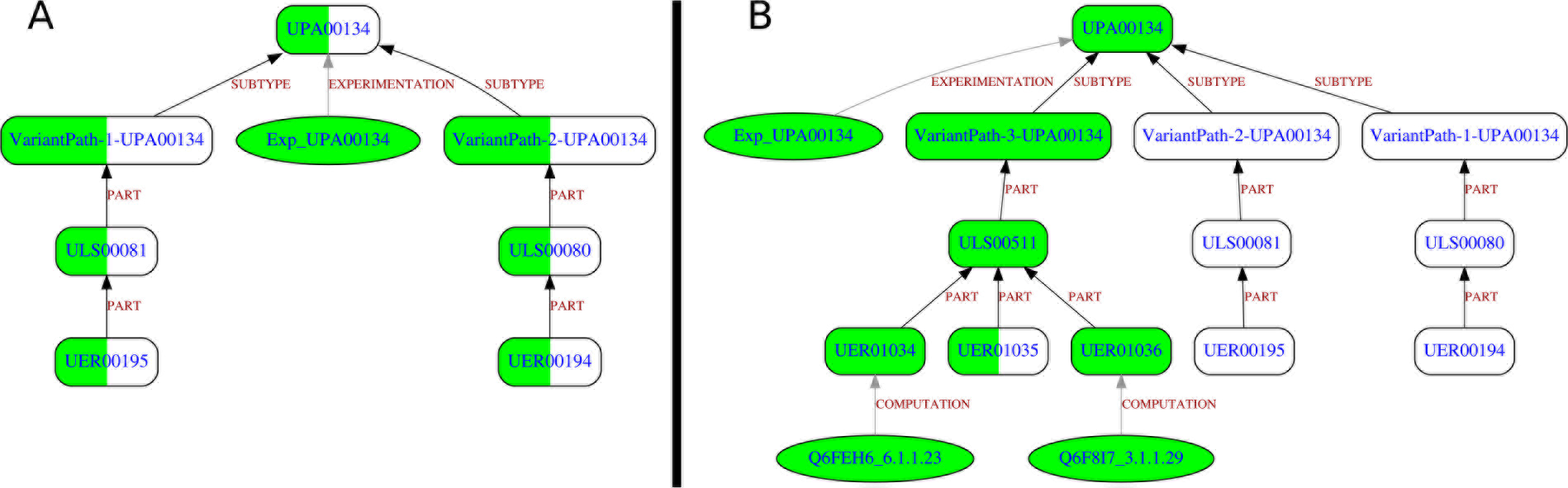
GROOLS results for asparagine biosynthesis in *Acinetobacter baylyi*. The asparagine biosynthesis in *Acinetobacter baylyi* was evaluated using UniPathway and UniProtKB. For GROOLS reasoning, the specific mode was used. Part A corresponds to the original pathway definition in UniPathway. Part B corresponds to the enhanced pathway definition with an additional variant. See Figure 3 for the legend and these links (http://www.genoscope.cns.fr/agc/grools/paper/fig4A.html, http://www.genoscope.cns.fr/agc/grools/paper/fig4B.html) for an interactive view.

### 3.3 UniPathway specific mode example

In UniPathway, two variants made of one enzymatic reaction are described for the biosynthesis of asparagine (UPA00134). Using *Acinetobacer baylyi* UniProtKB annotations (Figure 4 part A), any observation is linked to the two enzymatic reactions (UER00194 and UER00195) (i.e. there is any protein annotated with the corresponding UERs). Since this pathway is expected in *A. baylyi* (Exp_UPA00134 Observation), GROOLS reasoning concludes that both variants and reactions could be missing (i.e. GROOLS is not able to choose between those two). Unfortunately, UniPathway is wrong because a third pathway variant is known and reported in other metabolic databases like MetaCyc [19]. It corresponds to a tRNA-dependent transamidation mechanism for the conversion of aspartate to asparagine. This variant made of 3 reactions was integrated in the UniPathway model and the GROOLS reasoning was relaunched (Figure 4 part B). Using this modified model, GROOLS correctly selects the third variant by assigning a ‘Confirmed Presence’ conclusion and detects a missing reaction (UER01035). This latter is catalyzed in *A. baylyi* by an heterotrimer made of GatA, B, C proteins which cannot be assigned to the UER01035 as the corresponding EC number (6.3.5.6) is missing in their UniProtKB annotation.

## 4 Discussion

Following the initiative of Genome Properties [10] and IMG [9] systems, we present here a methodological framework for the evaluation of protein function predictions in the context of biological processes like metabolic pathways. Processes are modeled in a generic and hierarchical representation of knowledge, which is based on a conceptual graph with ‘part’ and ‘subtype’ relations. The reasoning is made of four rules that are activated dynamically by the propagation of observations over the graph. One original aspect of our work is the use of paraconsistent logic to deal with uncertainty and contradiction in a discriminating way. Indeed, predictions and expectations available for an organism cover only a fraction of biological processes. Moreover, these observations may contradict each other as they come from multiple resources with different levels of quality. At the end of the reasoning, GROOLS assigned 16 different conclusion values on all biological process components that represent confirmed, inconsistent, missing and other states. Genome Properties and IMG system, which rely on classical logic, are not able to make the distinction between conflicting and missing observations leading to rough conclusions. It should be also noticed that results of these methods cannot be easily compared to GROOLS ones as any software is available. GROOLS is distributed as an open source software and can be integrated at the end of various annotation workflows to evaluate functional annotations and pinpoint inconsistencies. It can analyze any annotated complete genome but requires that functional annotations could be linked to process components (e.g. enzymatic activities described by EC numbers and/or reaction identifiers). GROOLS was tested over 14 organisms with growth phenotype data. Analyses were made for two biological process resources (Genome Properties and UniPathway) and two sources of annotation (UniProtKB and MicroScope).

In terms of methodological limitations, GROOLS does not support quantitative values for observations (e.g. function prediction accuracy, optical density for growth phenotypes): they must be discretized in true and false values before the reasoning. Beside, GROOLS deals only with biological processes that can be described with specialization (‘subtype’) and composition (‘part’) relations, like metabolic pathways or macro-molecular systems (e.g. secretion systems). Thus, it is not designed to reason on processes containing regulation information (i.e. generally expressed with ‘positively or negatively regulate’ relations). Nevertheless, a new type of relations, named ‘avoid’, could be implemented in GROOLS. Unlike other relations between PK, ‘avoid’ relation will indicate concept components that should not be present. This type of relation could be useful to model more complex phenotypes like, for example, ‘strict anaerobe’ for which anaerobic metabolic pathways should be predicted, but not aerobic ones.

To increase the diversity of biological processes in GROOLS, we plan to integrate data from MetaCyc [19] that contains a comprehensive list of experimentally elucidated metabolic pathways with variant definition. Another resource of biological processes is the widely used Gene Ontology (GO) system [22]. It offers terms to describe functions and processes that are classified using ‘part’ and ‘subtype’ relations like in GROOLS model. Unfortunately, links between process and molecular function terms are missing in GO. For instance, it is difficult to retrieve the different sets of enzymatic activities that compose a metabolic pathway. This limitation is currently discussed in the GO consortium. They are moving to a new model called Logical Extension of the Gene Ontology (LEGO) that will serve as a basis for reasoning systems like the one presented in this paper.

To conclude, GROOLS, by its reactive nature and its fast execution time, could be integrated in an annotation system (such as MicroScope) as a live annotation companion to guide biologists during the curation process of their genome: at each protein annotation update, a new observation should be created in the GROOLS working memory and, then, would be propagated locally through the graph to alert users about impacts on biological process conclusions.

## 5 Conclusions

GROOLS software is designed to evaluate the overall accuracy of functional unit and pathway predictions according to organism experimental data like growth phenotypes. It allows users to compare different sources of annotations for a genome. Furthermore, using conclusion values, biocurators can quickly focused on missing or contradictory predictions to improve protein functional annotation. With the aim of building a metabolic model of an organism, and before applying time-intensive gap filling methods, GROOLS could thus be used to improve the initial metabolic network reconstruction in regards to the genome annotation and growth phenotype data.

## 6 Additional Files

### 6.1 Additional file 1 — Graphical illustration of GROOLS reasoning

### 6.2 Additional file 2 — Accuracy with detailed counts

## 7 Acknowledgements

We would like to thank Anne Morgat for her valuable advice about UniPathway data, Thierry Lombardot for his help in SPARQL UniProtKB queries and Alain Viari for his support and constructive comments.

## 9 Availability of data and materials

GROOLS is a free open source software distributed under CeCILL license and can be downloaded from https://github.com/Grools. Detailed results are available at https://www.genoscope.cns.fr/agc/grools.

## 10 Author’s contributions

DV and CM supervised the work. JM and DV designed the method and conducted the analyses. JM implemented the software. AJ conducted developments for data extraction and gave support to manuscript preparation. JM and DV wrote the manuscript.

## 12 Consent for publication

Not Applicable.

## 13 Ethics approval and consent to participate

Not Applicable

